# Functional specialization of the hippocampus along its dorso-ventral axis: from span to supraspan in working memory capacity

**DOI:** 10.1101/2025.10.07.680879

**Authors:** Filomena Grazia Alvino, Anna Carboncino, Diego Carrella, Jenny Crain, Attilio Iemolo, Nadia Concetta Giordano, Laura Olivito, Diletta Cavezza, Maria De Risi, Diego Di Bernardo, Alessandro Treves, Elvira De Leonibus

**Affiliations:** Telethon Institute of Genetics and Medicine (TIGEM), Via Campi Flegrei 34, 80078 Pozzuoli, Napoli, Italy; Institute of Biochemistry and Cell Biology (IBBC), CNR, Via Ercole Ramarini 32, 00015 Monterotondo, Rome, Italy; SISSA – Cognitive Neuroscience, via Bonomea 265, Trieste, Italy

**Keywords:** Hippocampus, Working memory capacity, Dorsal and Ventral regions, Spatial memory, allocentric, egocentric navigation

## Abstract

Working memory (WM) is characterized by a limited capacity or *span*, beyond which performance deteriorates, the *supraspan* condition. Human neuropsychology and neuroimaging studies have implicated the medial temporal lobe (MTL), including the hippocampus (HPC), in WM particularly when memory load exceeds span, yet no animal model has been established to investigate supraspan mechanisms. Here, we present a novel rodent model of supraspan memory and examine the distinct roles of dorsal and ventral HPC. Using selective NMDA lesions in mice combined with object recognition tasks and a modified radial arm maze (RAM), we manipulated memory load and strategy constraints to assess span and supraspan performance. Dorsal HPC lesions impaired object memory span, while ventral lesions spared this ability. In the RAM, span performance (number of correct arm visits before the first error) was unaffected by either lesion under allocentric conditions. However, once animals exceeded span, following the first error, here defined as the supraspan condition, both dorsal and ventral lesions caused catastrophic disorientation and error escalation. Under free-choice conditions, ventral HPC lesions selectively reduced the occurrence of trials with no errors in the first 5 choices, increased errors overall, and disrupted sequential egocentric strategies. Together, these results demonstrate that supraspan performance can be modeled in rodents, revealing a dorsal contribution to object memory under high load, a ventral contribution to egocentric spatial strategies, and a shared hippocampal role in supraspan memory performance. This paradigm provides a translational framework for investigating supraspan deficits observed in MTL patients and Alzheimer’s disease.

## Introduction

Broadly defined, working memory (WM) is a temporary buffer that maintains information in service of ongoing behavior. A hallmark of WM is its limited capacity, or *span*, operationally quantified by the (average) maximum number of items that can be retained in immediate memory^1^. Tasks in which the load remains within this capacity are termed *subspan*, while *supraspan* tasks exceed it, requiring additional mechanisms beyond immediate WM.

In humans, neuropsychological evidence has demonstrated that medial temporal lobe (MTL) lesions impair performance specifically under supraspan conditions. The classic case of H.M., who retained information briefly if undistracted but could not sustain it once capacity was exceeded, first suggested this dissociation^2^. More recent studies confirmed that amnesic patients with selective hippocampal lesions perform normally on span tasks but fail when memory load surpasses span, even at short delays^3,4^. Deficits in supraspan list-learning tests, such as the Rey Auditory Verbal Learning Test, correlate with hippocampal pathology in epilepsy^5^ and predict progression in Alzheimer’s disease. Functional imaging has corroborated these findings. Hippocampal and subicular activation scales with memory load during item recognition and n-back tasks^6–8^, and persistent activity in the human HPC during WM predicts subsequent long-term memory formation^9^. Thus, supraspan performance reveals hippocampal contributions not evident under subspan conditions.

Despite this evidence, no validated animal model of supraspan memory exists. Rodent studies have documented WM capacity limits^10–12^ and fronto-striatal circuits have been implicated in span regulation^13–15^. Using a modified version of an incidental object recognition paradigm, in which the memory load is increased by increasing the number of objects to which the animals are exposed during the study phase (the different objects task/Identical objects task-DOT/IOT) it was shown that hippocampal lesions restricted to the dorsal portion of the HPC impairs recognition when animals are exposed to 6, but not to 4, different objects^11,16^. While the DOT/IOT allows to test incidental memory in conditions of overload^15^, it does not allow to exactly distinguish between span and superspan conditions, within each single trial. Additionally, the HPC is not a homogeneous structure; rather, an anatomical distinction between the dorsal and the most ventral part of the HPC, along the septo-temporal axis, has been demonstrated^17^. Dorso-ventral subregions originate from separate genetic programs and are integrated into distinct anatomical and cellular circuits^18–22^ . They also appear to be functionally distinct, with the dorsal HPC primarily associated with spatial memory^23–26^, while the ventral HPC is more closely linked to egocentric spatial memory and emotional behaviour^27–31^. Some reports describe speared span after complete HPC lesions^10^, while others emphasize the dorsal HPC in spatial WM^25,26^ and the ventral HPC in emotional regulation and egocentric strategies^28–31^. Whether this dorso-ventral specialization extends to WM capacity, and particularly to supraspan performance, has not been tested.

Here, we address this gap. We define supraspan in rodents as the condition following the first error in a radial arm maze task, when we reckon that immediate WM resources have been exceeded. By combining object recognition (6-DOT/6-IOT) and a modified radial arm maze under allocentric and egocentric strategy conditions, with selective lesions of dorsal and ventral HPC, we provide the first evidence that supraspan can be modeled in rodents. We further demonstrate that dorsal and ventral HPC play distinct but complementary roles in these forms of working memory, with dorsal HPC supporting high-load object WM, ventral HPC supporting egocentric spatial strategies, and both subregions required for supraspan maintenance in the radial arm maze.

## Results

### Dorsal Hippocampus, but not Ventral Hippocampus, regulates object memory capacity in mice

To investigate the functional dissociation along the dorso-ventral axis of the HPC in regulating memory capacity, we performed selective ablation by administering high concentrations of NMDA using different sets of stereotaxic coordinates that could differentiate the two regions along the dorso-ventral / septo-temporal axis. The extent of ablation ranged from localized damage confined to the CA hippocampal fields to broader lesions encompassing the entire subregion either the dorsal or ventral HPC (Fig 1A-B). These ablation procedures thus effectively dissociated two subregions, here defined as Dorsal HPC and Ventral HPC, with distinct anatomical connections, which were assessed using Fluorogold injections in each subregion. The rhomboid thalamic nucleus and the reuniens thalamic nucleus were confirmed to be selectively connected with the dorsal HPC^32^. In mice injected in the ventral HPC, a higher signal intensity was observed in the stria terminalis, the lateral hypothalamus, and the posterior part of the basolateral amygdaloid nucleus^33,34^ (Fig 1C).

**Figure 1.**
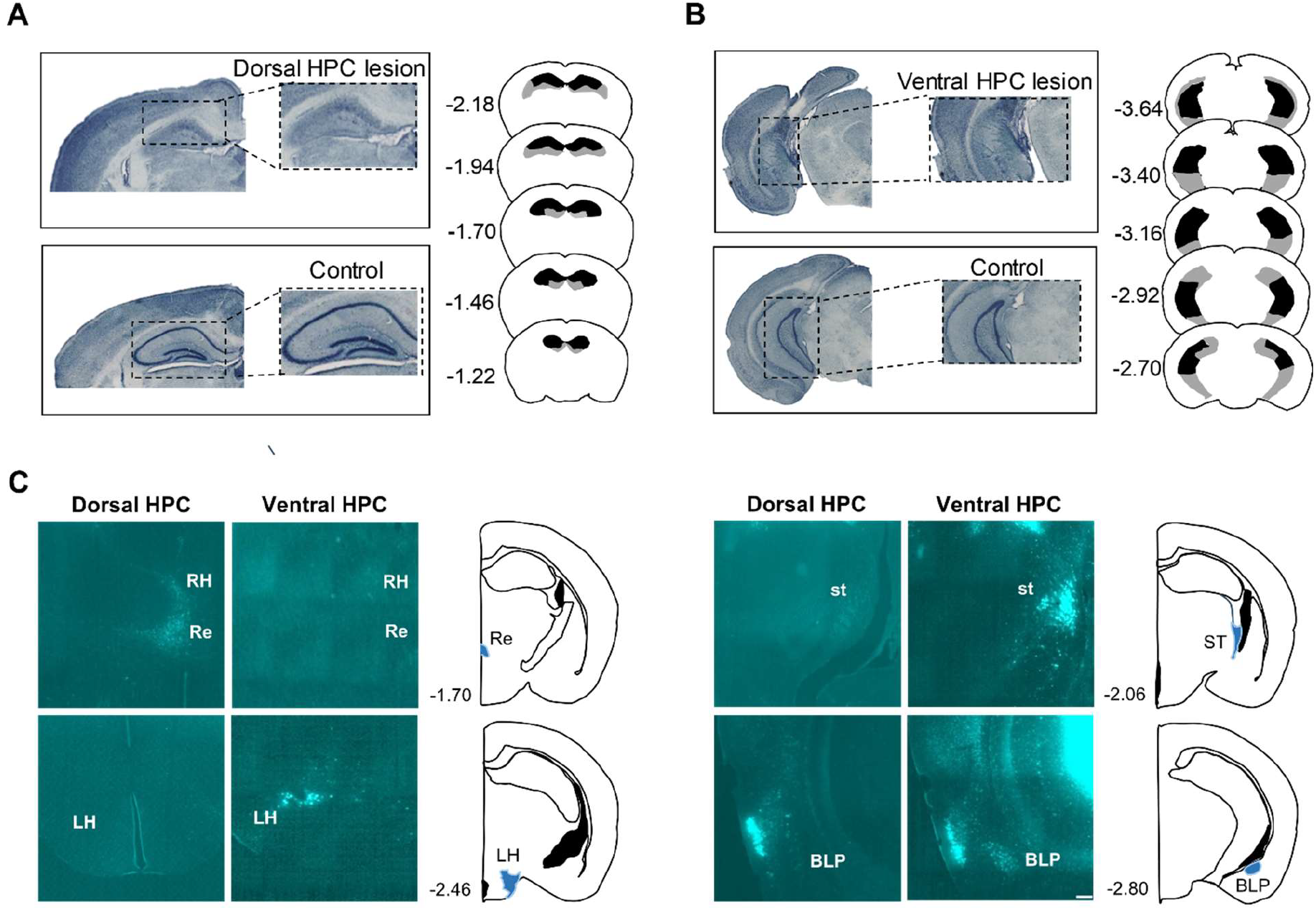
Anatomical dissociation between the Dorsal and Ventral Hippocampal subregions. **A)** Photomicrographs of Nissl-stained coronal sections illustrating the dorsal hippocampus (HPC) lesion compared to the Control group (left). A graphic reconstruction (right) shows the largest (grey areas) and smallest (black areas) extents of the lesion in the dorsal lesion group (right). **B)** Photomicrographs of Nissl-stained coronal sections illustrating the ventral HPC lesion compared to the Control group (left). A graphic reconstruction (right) shows the largest (grey areas) and smallest (black areas) extents of the lesion in the ventral lesion group (right). The numerical values adjacent to the slices indicate the distance in millimeters relative to bregma in the anterior-posterior plane^35^. **C)** Fluorogold mediated labeling of the afferent connections to the dorsal HPC and ventral HPC shows the differential anatomical connectivity of the two subregions. Rhomboid thalamic nucleus (RH) and reuniens thalamic nucleus (Re) are selectively connected with the dorsal HPC. On contrary, selective labeling of the stria terminalis (st), lateral hypothalamus (LH) and posterior part of the basolateral amygdaloid nucleus (BLP) is seen upon ventral but not dorsal HPC fluorogold injection.

To investigate the functional dissociation between the two subregions, the injected mice underwent a battery of behavioral tasks following surgery (Fig 2A). First, we demonstrated that the Ventral, but not the Dorsal, hippocampus plays a role in anxiety-like behavior, as previously documented^29,30^. Specifically, lesions in the ventral HPC increased both the time spent and the number of entries into the open arms of the elevated plus maze task, in agreement with previous experimental evidence reporting a preferential role of the ventral HPC in regulating anxiety-related behaviors (Elevated plus maze % open arms time: one-way ANOVA, lesion F _(2,29)_ = 4.650; p = 0.018; *Tukey’s multiple comparisons test*: Control vs. Dorsal HPC lesion p = 0.587, Control vs. Ventral HPC lesion p = 0.016, % open arms entries: F _(2,29)_ = 4.472; p = 0.020; *Tukey’s multiple comparisons test*: Control vs. Dorsal HPC lesion p = 0.99, Control vs. Ventral HPC lesion p = 0.048). (Fig 2B).

**Figure 2.**
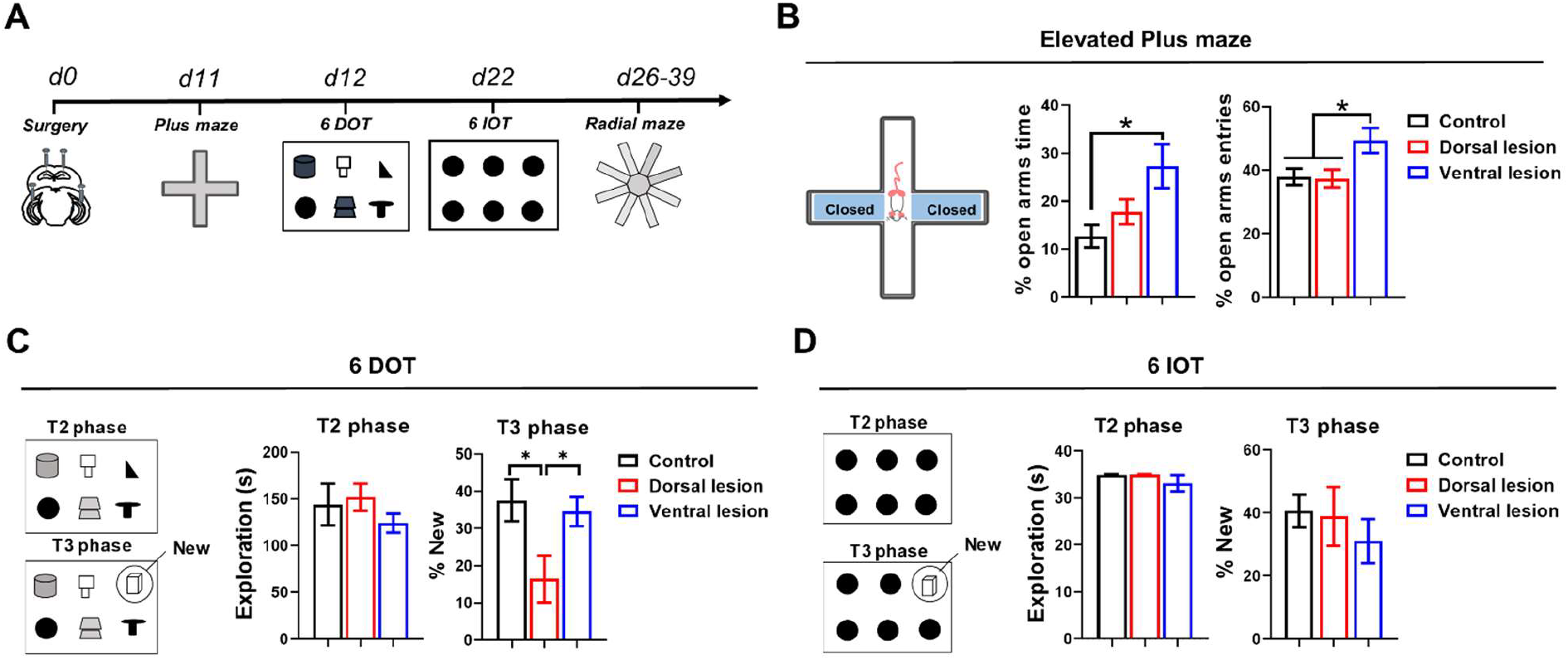
Dorsal and Ventral HPC roles in object memory and in anxiety. **A**) Experimental timeline of the behavioral tasks following surgical ablation of the dorsal and ventral hippocampus (HPC) subregions. **B)** Ventral but not Dorsal HPC lesions reduced anxiety-like behavior as assessed in the elevated plus maze (% open arms time: one-way ANOVA, lesion F _(2,29)_ = 4.650; p = 0.018; Tukey’s multiple comparisons test: Control vs. Ventral HPC lesion p = 0.016, % open arms entries: F _(2,29)_ = 4.472; p = 0.020; Tukey’s multiple comparisons test: Control vs. Ventral HPC lesion p = 0.048).**C)** Schematic representation of the T2 and T3 phases of the 6-DOT task, with the new object in the T3 phase circled (left). Total object exploration during the T2 phase is not affected by either HPC lesion (one-way ANOVA, lesion F _(2,25)_ = 0.957, p=0.398) (middle). However, the dorsal lesion group shows a reduced percentage of new object exploration compared to both Control and Ventral lesion group (one-way ANOVA, lesion F _(2, 25)_ = 4.520; p = 0.0211, Tukey’s multiple comparisons test: Control vs. Dorsal HPC lesion p = 0.030) (right). **D)** Schematic representation of the T2 and T3 phases of the 6-IOT task, with the new object in the T3 phase circled (left). Total object exploration during the T2 phase is not affected by either HPC lesion (one-way ANOVA, F _(2,25)_ =0.686; p = 0.513) (middle). The percentage of new object exploration relative to total exploration remains unimpaired following HPC lesions (one-way ANOVA, lesion F _(2,25)_ =0.5097; p = 0.607) (right). Data are expressed as mean ± SEM. * p < 0.05 Turkey’s multiple comparisons test.

To assess the capacity of one form of object memory, we utilized the 6-Different Objects Task (6-DOT) and the 6-Identical Objects Task (6-IOT), which evaluate spontaneous object recognition memory under conditions of high (6-DOT) and low (6-IOT) memory load^11,16,36–38^. Previously, we demonstrated that a dorsal HPC lesion reduces such object memory capacity to four different objects, compared to the normal limit of six different objects observed in control mice^11^. In the present study, we confirmed that a dorsal HPC lesion impairs novel object recognition in the 6-DOT condition, whereas a ventral HPC lesion does not (T3 phase: one-way ANOVA, lesion F _(2,25)_ = 4.520; p = 0.0211; *Tukey’s multiple comparisons test*: Control vs. Dorsal HPC lesion p = 0.030, Control vs. Ventral HPC lesion p = 0.902) (Fig. 2C). However, neither type of lesion impairs performance in the 6-IOT condition (T3 phase: one-way ANOVA, lesion F _(2,25)_ =0.5097; p = 0.607 (Fig. 2D). We also ruled out the possibility that a general loss of interest in novelty or a compromised exploration pattern influenced this finding, as neither lesion affected object exploration (T2 phase 6 DOT: one-way ANOVA, lesion F _(2,25)_ = 0.957, p=0.398; 6-IOT: F _(2,25)_ =0.686; p = 0.513) or locomotor activity (T1 phase 6-DOT: lesion F _(2,25)_ = 1.571, p = 0.228; 6 IOT: lesion F _(2,25)_ = 1.214, p = 0.314) (Fig S1).

Taken together, these findings suggest a functional dissociation between the ventral and dorsal HPC subregions in regulating memory capacity for object-related information.

### Differential involvement of Dorsal and Ventral hippocampus in allocentric and egocentric spatial memory span and supraspan

To assess spatial WM span, we used a modified version of the radial arm maze, manipulating memory loadby allowing animals to retrieve food from 3, 6, or 8 open arms^16^. In our design, the test was initially conducted under a Confinement procedure (first 4 days), which encouraged the use of allocentric strategies by restricting the use of egocentric ones (Fig. 3A). In the following 4 days, the confinement was removed, allowing the animals to freely adopt any strategy to solve the task (No Confinement) (Fig. 3C).

**Figure 3.**
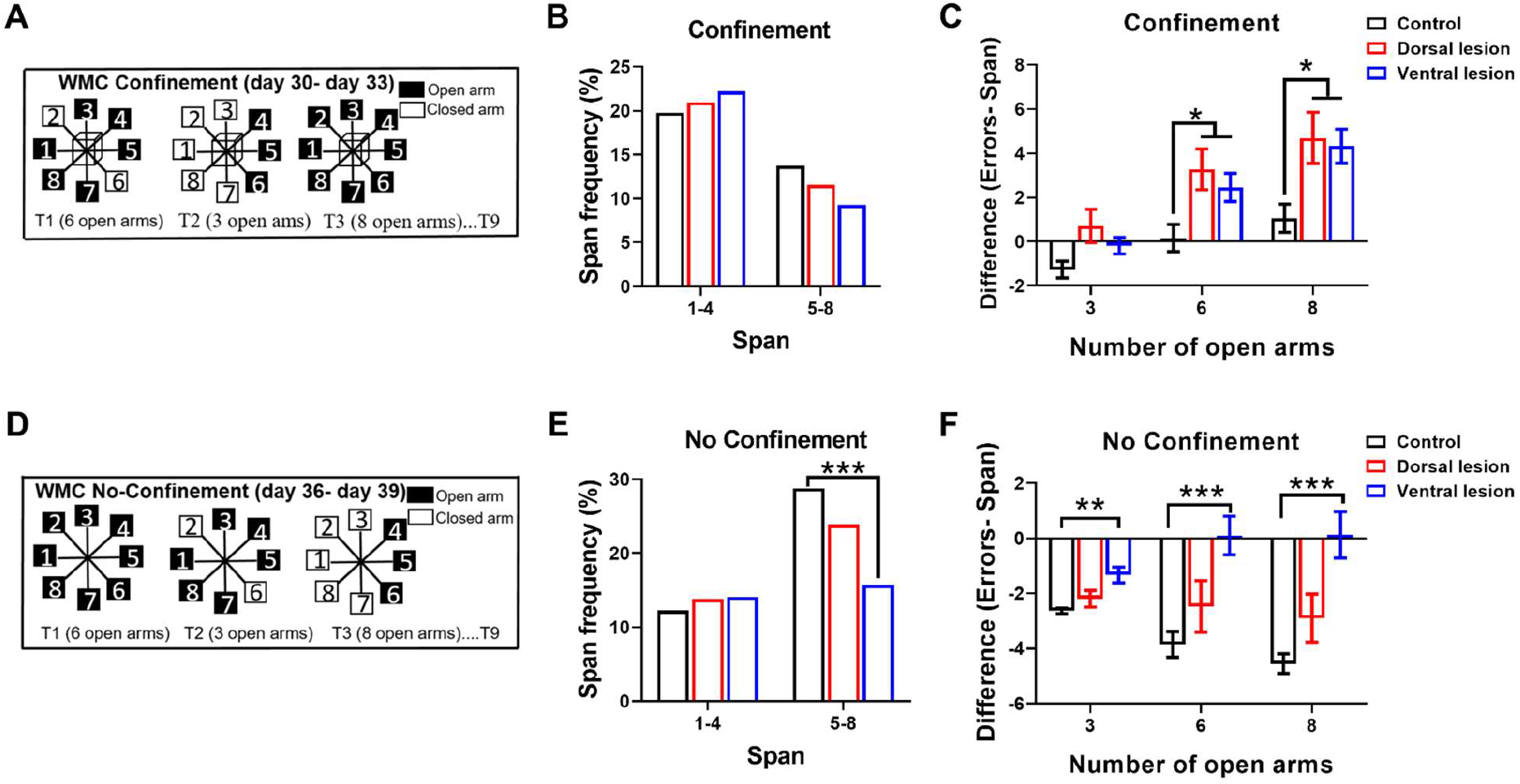
Dorsal and Ventral HPC role in spatial working memory capacity. **A)** Schematic representation of the working memory capacity training phase under the Confinement procedure. In this phase, the working memory (WM) load was manipulated by varying the number of open arms between 3, 6, and 8 (black squares). The three conditions were randomly alternated between trials, with the specific open arms being randomly selected within trials and across days. **B)** The frequency of lower spans (Span 1–4) and higher spans (Span 5–8) was not significantly affected by either lesion groups (Fischer’s exact test with Hochberg-Benjamini multiple testing correction).**C)** In the Confinement procedure, both dorsal and ventral HPC lesion affected the supraspan index (Errors – Span), in the 6 and 8 open arms conditions (F _(2,29)_ = 5.924, p=0.0069, with Tukey’s multiple comparisons test) Data are expressed as mean ± SEM. **D)** Schematic representation of the WMC training phase under the No Confinement procedure. The number and configuration of the open arms (black squares) were randomly changed across trials and days. **E)** The frequency of higher spans (Span 5–8) was significantly reduced in the ventral HPC lesion group (Fischer’s exact test with Hochberg-Benjamini multiple testing correction). **F)** The supraspan index was affected upon ventral HPC lesion, but not the dorsal HPC lesion, across all memory load conditions (two-way ANOVA, lesion x number of open arms F _(4, 58)_ = 5.769, p=0.0006 with Tukey’s multiple comparisons test) Data are expressed as mean ± SEM. **F)** The frequency of higher spans (Span 5–8) was significantly reduced in the ventral HPC lesion group (Fischer’s exact test with Hochberg-Benjamini multiple testing correction). * p < 0.05, ** p < 0.01, *** p < 0.001.

Neither type of lesion impaired the learning of the general rules of the task (Pre-training phase: one-way ANOVA, lesion F _(2,29)_ = 2.262, p=0.12) (Fig S2A). To assess WMC in the Confinement procedure, we evaluated the Span, defined as the number of arm visits before committing the first error. We observed that the occurrence of lower (Span 1-4) and higher (Span 5-8) spans did not differ significantly between the groups (Fischer’s exact, p = 0.05 *Hochberg-Benjamini multiple testing correction*) (Fig 3B). Conversely, when we measured the mean number of arm re-entries (errors), a significant effect of the lesion was observed (two-way repeated measures (RM) ANOVA, lesion F _(2,29)_ = 6.843, p=0.0037) which did not appear to be strongly dependent on memory load (lesion x number of open arms, F _(4, 58)_ = 1.331, p=0.269 (Fig S2B). These findings suggest that, regardless of memory load, HPC lesions impaired the ability to solve the task, leading to disorientation. We calculated a supraspan index by computing the difference between the mean number of errors and the mean span for each animal. This index, combining span impairment with supraspan errors was significantly increased after HPC lesions (two-way RM ANOVA, lesion F _(2,29)_ = 5.924, p=0.0069; (*Tukey’s multiple comparisons test*, 6 open arms: Dorsal lesion vs. Control p = 0.032, Ventral lesion vs. Control p = 0.043; 8 open arms: Dorsal lesion vs. Control p = 0.039, Ventral lesion vs. Control p = 0.010) (Fig. 3C). Indeed, while the values in the control group were around 0, for the lesioned groups they were around 4-5, implying that they visited several additional arms before completing the task.

These findings suggest that neither dorsal nor ventral HPC lesions have a detrimental effect on memory span in a spatial WMC version of the radial arm maze task when allocentric strategies are enforced. Nevertheless, both HPC lesions caused an increased number of errors after the first one was committed, suggesting impaired post-error WM maintenance.

On the other hand, when the WMC radial arm maze task was administered under the No Confinement procedure, which allowed for the choice of any strategy to solve the task, Ventral but not Dorsal HPC lesion produced a significant decrease in the occurrence of higher span lengths, ranging from 5 to 8 (Fischer’s exact, *Hochberg-Benjamini multiple testing correction* Ventral lesion vs. Control p < 0.001) (Fig 3E). Additionally, the mean number of errors was robustly affected by ventral, but not dorsal, HPC lesions across all three memory load conditions (lesion x number of open arms F _(4, 58)_ = 4.761, p=0.002) (Fig S2C). Similarly, both the control and dorsal HPC lesion groups showed negative values of the “errors-span” index, again indicative of a low number of errors in this condition, whereas the ventral HPC lesion led to approximately null values (lesion x number of open arms F _(4, 58)_ = 5.769, p=0.0006; Tukey’s multiple comparisons test, 3 open arms: Ventral lesion vs. Control p = 0.002, 6 open arms: Ventral lesion vs. Control p = 0.0005, 8 open arms: Ventral lesion vs. Control p = 0.0004) (Fig.3F).

Together, these findings suggest that dorsal and ventral HPC are not differentially engaged in the spatial WMC task when an allocentric spatial strategy is favored. Both HPC lesions more severely impaired the cognitive processes involved supraspan than in memory span. Instead, the ventral HPC appears to play a critical role when the spatial strategy for solving the task can be freely chosen.

### The Ventral Hippocampus is more involved in the acquisition of an egocentric sequential strategy

Prompted by our findings suggesting a crucial role of the Ventral HPC in the execution of the strategy to solve the task, we next sought to investigate whether the dorsal and ventral HPC play a role in the choice patterns of arms. Previous studies have demonstrated that, in a No Confinement procedure of the radial arm maze task, control rats preferentially employ sequences of 45° turns corresponding to a serial clockwise or counter-clockwise strategy starting from the first three or four trials. In contrast, hippocampal-lesioned animals show impairments in this strategy^39–41^. However, when a Confinement procedure is introduced, rats do not exhibit a particular preference for any strategy^42,43^.

To test this in our paradigm, we calculated the scores for adjacent, sequential, alternating, opposite, or random strategies using the custom-built, open-source database *RAM* (Fig 4A-B). In the Confinement procedure, both control and lesioned animals exhibit similar choice patterns with both 6 (two-way repeated measures (RM) ANOVA, lesion F _(2, 29)_ = 0.2977, p=0.744) and 8 open arms (lesion F _(2, 29)_ = 1.202, p=0.315) (Fig 4C). Conversely, in the No Confinement procedure, ventral HPC lesion robustly decrease the use of the adjacent strategy with both 6 (lesion x strategies F _(6, 87)_ = 7.029, p < 0.0001; *Tukey’s multiple comparisons test*, Adjacent: Ventral lesion vs. Control p = 0.0008) and 8 open arms (lesion x strategies F _(6, 87)_ = 6.481, p < 0.0001; *Tukey’s multiple comparisons test*, Adjacent: Ventral lesion vs. Control p = 0.0006) (Fig 4D).

**Figure 4.**
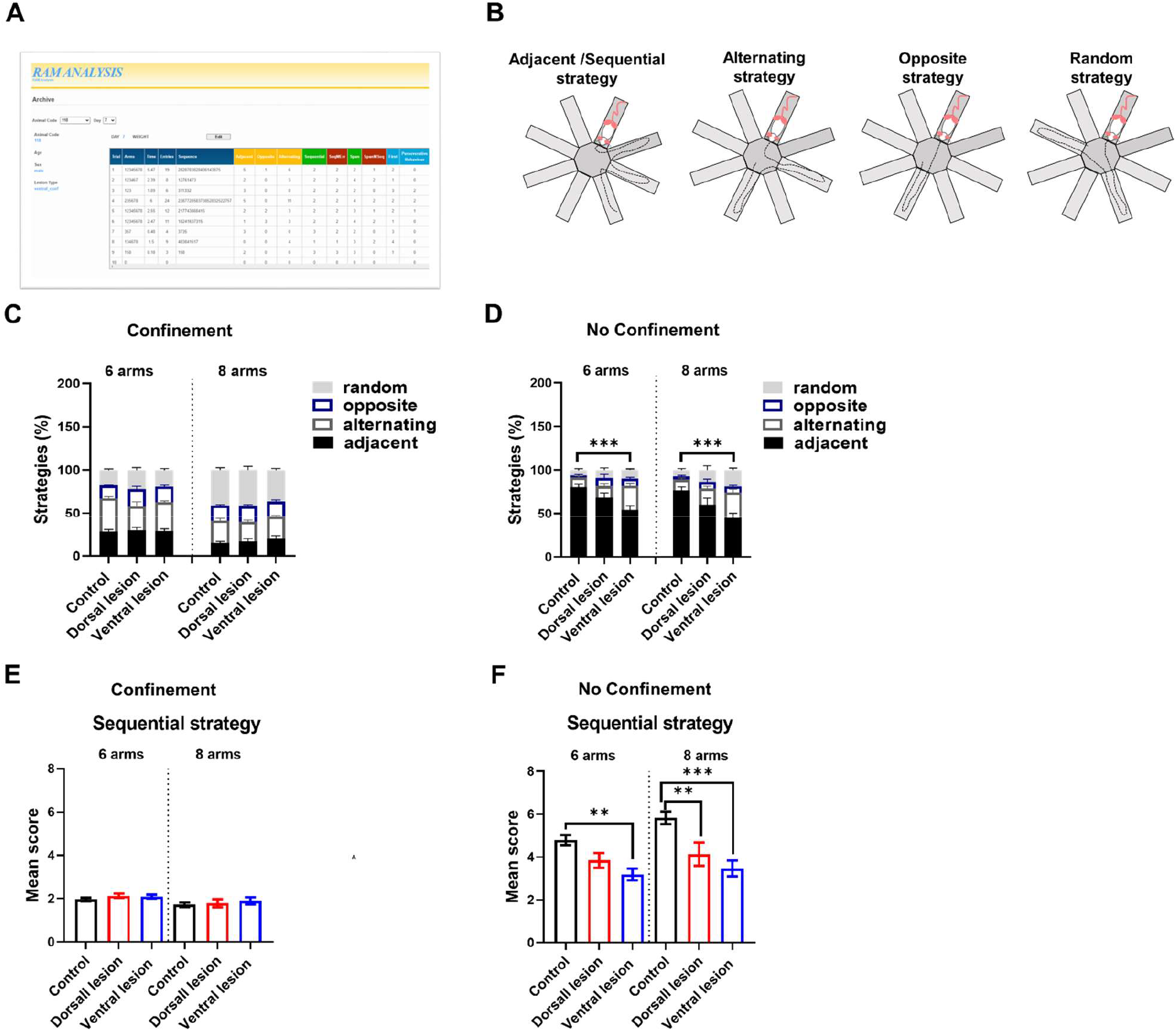
Ventral and Dorsal HPC involvement in the acquisition of an egocentric sequential strategy. **A)** Example of a web page of the RAM analysis database used for the automatic scoring of the strategies used in the RAM task. **B)** Schematic representation of the four possible strategies used in the task (Adjacent/sequential, alternating, opposite and random). **C)** Percentage of choices made in adjacent, alternating, opposite or random arms in the Confinement procedure. Neither HPC lesion affects the pattern of choices made in either 6 open arms condition (two-way repeated measures (RM) ANOVA, lesion F _(2, 29)_ = 0.2977, p=0.744) or 8 open arms condition (lesion F _(2, 29)_ = 1.202, p=0.315). **D)** Percentage of choices made in adjacent, alternating, opposite or random arms in the No Confinement procedure. Ventral but not Dorsal HPC lesions significantly affect the choices pattern made in the task, reducing the use of adjacent choices with both 6 open arms (lesion x strategies F _(6, 87)_ = 7.029, p < 0.0001; Tukey’s multiple comparisons test, Adjacent: Ventral lesion vs. Control p = 0.0008) and 8 open arms (lesion x strategies F _(6, 87)_ = 6.481, p < 0.0001; Tukey’s multiple comparisons test, Adjacent: Ventral lesion vs. Control p = 0.0006). **E)** Neither Dorsal nor Ventral HPC lesions affected the use of the Sequential strategy in the Confinement procedure in 6 and 8 open arms conditions (two-way RM ANOVA, lesion F _(2, 29)_ = 0.604, p = 0.553). **F)** Ventral, more than the Dorsal HPC impaired the use of the egocentric sequential strategy with both 6 open arms and 8 open arms (two-way RM ANOVA, lesion F _(2, 29)_ = 8. 931, p = 0.001 followed by Sidak’s multiple comparisons test). Data are expressed as mean ± SEM. * p < 0.05, ** p < 0.01, *** p< 0.001.

Guided by this finding, we also assessed the score of the sequential egocentric strategy, a measure of the longest sequence of correct adjacent arm choices. As previously reported, the Confinement procedure reduced the use of this strategy, maintaining an overall average mean length of 2.063 in the 6 open arms condition and 1.799 in the 8 open arms condition, with no evidence of any impact of the lesion on this effect (two-way RM ANOVA, lesion F _(2, 29)_ = 0.604, p = 0.553) (Fig. 4E). Conversely, in the No Confinement procedure, the lesion had a robust effect on the use of the strategy (two-way RM ANOVA, lesion F _(2, 29)_ = 8. 931, p = 0.001), with the ventral HPC significantly decreasing it in both the 6 (*Sidak’s multiple comparisons test*, Ventral lesion vs. Control p = 0.006) and 8 open arms conditions (*Sidak’s multiple comparisons test*, Ventral lesion vs. Control p < 0.0001), while the dorsal HPC reduced it significantly only when the number of open arms was 8 (*Sidak’s multiple comparisons test*, Dorsal lesion vs. Control p = 0.0053) (Fig 4F).

Notably, when the learning process of the task was analysed across days under both the Confinement and No Confinement procedures, a differential engagement of the two subregions was also evident. In the Confinement procedure, a mild Lesion × Days interaction effect was observed (F _(6, 87)_ = 2.303, p = 0.041), with the ventral HPC lesion appearing to facilitate the learning process by reducing the mean number of errors across days only in the 8 open arms condition (*Tukey’s multiple comparisons test*, day 4 vs. day 2 Ventral lesion vs. Control p = 0.0092) (Fig. S3A). Interestingly, when switching from the Confinement to the No Confinement procedure, the dorsal HPC-lesioned animals improved their performance, whereas the ventral HPC-lesioned animals did not (*Tukey’s multiple comparisons test*, 6 open arms: Dorsal lesion day 4 No Confinement vs. day 4 Confinement p =0.04; 8 open arms: Dorsal lesion day 1 No Confinement vs. day 4 Confinement p =0.031) (Fig S3B).

In parallel, a learning process of the sequential strategy was not observed in the Confinement procedure (Fig S3C). However, upon switching from the Confinement to the No Confinement procedure, a lesion × days interaction was observed (6 open arms: lesion x days F _(8, 116)_ = 2.099, p = 0.041, 8 open arms: lesion x days F _(8, 116)_ = 3.359, p = 0.002), with the ventral HPC lesion leading to a significant difference from both Control and Dorsal groups (6 open arms: *Tukey’s multiple comparisons test*, day 1 Ventral lesion vs. Control p = 0.011, day 2 Ventral lesion vs. Control p = 0.009, day 3 Ventral lesion vs. Control p = 0.005; 8 open arms: day 1 Ventral lesion vs. Control p = 0.037, day 3 Ventral lesion vs. Control p = 0.001, day 4 Ventral lesion vs. Control p = 0.0005) (Fig S3D).

## Discussion

In this study, we investigated the functional specialization of the HPC along its dorso-ventral (septotemporal) axis in regulating memory capacity. Our findings demonstrate a dorsal specialization in incidental object memory and a ventral one in egocentric spatial working memory and anxiety. In contrast, reaching the normal spatial span does not necessitate the activation of the HPC; however, upon the very first error, lesioned animals get disoriented and accumulate several errors, re-entering already visited arms, independently of the lesion site. This behavior suggests a selective “supraspan” deficit. The same radial maze protocol, where the use of egocentric strategies is allowed, highlighted the prominent role of the ventral HPC. The No-Confinement procedure strongly favored an egocentric sequential strategy, with both subregions involved, though the ventral HPC played the major role. To our knowledge, this is the first study to examine the role of HPC, and the functional specialization of its subregions, in span and supraspan memory.

Previous research examining the HPC role in object recognition memory in rats have focused on its involvement in delay-dependent object recognition memory. Rats with HP lesions displayed impaired object recognition memory when tested with delay intervals longer than 1 minute^44,45^. These findings suggest that shorter delays allow the task to be solved by simply judging the familiarity of the previously encountered stimulus, a process that could be fully mediated by sensory and/or association neocortex regions, without requiring involvement of the HPC^46^. Later studies demonstrated that the dorsal HPC is not involved in novelty detection with only two objects when the lesion is focal^47,48^. However, lesions covering 75-100% of the total dorsal HPC volume impaired memory for distinguishing two different objects after longer retention intervals^49^.

In a previous study, we demonstrated that a dorsal HPC lesion (75% of total hippocampal tissue) impaired novel object recognition memory within a short time interval (1 minute) only when six objects had to be recognized, but not with three or four objects. This was the first study to report the involvement of the mouse dorsal HPC in incidental object memory under high memory load conditions^11^. Consistent with this, a role for the HPC in object recognition under high memory load conditions has also been observed in humans^3^ and non-human primates^50^. In a visual short-term memory capacity task conducted during fMRI, patients exhibited a positive relationship between their WMC performance and the intensity of the signal in the bilateral posterior HPC^51^. Moreover, fMRI scans of subjects performing an operation span task and an arithmetic span task showed significant bilateral activation of the HPC, with greater involvement of the posterior region^52^. Here we expanded on these previous observations and showed that a dorsal HPC lesion impaired recognition of six different objects, while animals with a ventral HPC lesion did not show this impairment. The observed differences between the dorsal HPC and ventral HPC in object recognition memory is consistent with its preferential connectivity and functional commonalities with perirhinal and lateral entorhinal input^53–56^.

While the DOT/IOT allows to manipulate memory capacity by increasing the number of unfamiliar objects and serves as a model of incidental learning, it does not allow to evaluate whether suspraspan mechanisms are engaged in the task. Here, we investigated the role of the dorsal and ventral HPC in a radial arm maze procedure designed to assess three different memory load conditions (3, 6, and 8 baited open arms) and various spatial memory strategies. Animals were exposed daily to a random sequence of 3, 6, and 8 open and baited arms, with the order of trials and the configuration of open arms being randomly varied. This protocol was carried out over 4 days with a confinement procedure, followed by 4 days of a no-confinement procedure.

Our definition of supraspan performance aligns with that used in clinical studies. Jeneson and Squire^4^ proposed that MTL lesions impair performance only when immediate memory is insufficient, i.e., under supraspan conditions. Similarly, in our radial maze, control animals could recover after an error, while lesioned animals became disoriented. We interpret the error as a distractor, requiring reactivation of schemata or “intermediate buffers” to sustain ongoing performance. The hippocampus may help provide these backup mechanisms, consistent with proposals that it supports schema reactivation^57^ and error-related adjustments ^58,59^. Our results therefore demonstrate experimentally in rodents what has been inferred in patients: supraspan performance is hippocampal-dependent.

One of the most interesting observations arising from the results is that, in the Confinement procedure, neither of the two lesions reduced the span, as measured by the number of arms visited before committing the first error. This suggests that a single HPC subregion may be sufficient to preserve spatial WMC. Previous studies, in fact, have reported WMC impairments only in cases of complete HPC lesions^10^. However, after the first error, while Control animals can manage to complete the task with only one or two additional arm visits, lesioned animals become disoriented and accumulate a significant number of re-entries. The most conservative explanation for this finding is that, despite similar span in both control and lesioned groups, they rely on different memory strategies. Jeneson and Squire^4^ proposed that medial temporal lobe (MTL) lesions impair performance only when immediate and working memory are insufficient to support task demands— that is, under *supraspan* conditions. The deficit of HPC lesioned animals emerges immediately after the first error, a situation that may reflect a diversion of attention. Jeneson and Squire argue, however, that in such cases performance relies on long-term memory, even when the retention interval is brief. In the radial maze, each trial is unique and the occurrence of an error is unpredictable, making it difficult to specify whether long-term memory mechanisms are engaged. We speculate that errors function as ‘noise’ or ‘distractors’ in WM performance. The impact of this noise on memory might in part be mitigated by resorting to behavioural schemata pre-established in long-term memory, to the reactivation of which the HPC contributes^57^. Consequently, the pattern of errors observed in HPC lesioned groups may reflect an impairment in the reactivation of such schemata^58^. In other words, our HPC lesions could affect the post-error “back-up options” through which individuals resolve the conflict generated by the error and mobilize cognitive and attentional resources to pursue their goals^59^. Although this increase in errors following both types of HPC lesion appears to contradict previous evidence suggesting a specific role for the dorsal HPC^26^, it is important to consider that our protocol alternates open arm configurations across trials and days, rather than using a repeated full baited 8-arms task where the setting is always the same. Note that we find that across days, with 8 open arms, so that the configuration remains constant, a ventral HPC lesion actually leads to an improved performance. This suggests that a repeated 8-arm configuration could be less sensitive, after errors, to the disrupted activation of remedial schemata, following ventral HPC lesions.

Next, we found that during the transition to the No Confinement procedure, the ventral HPC, but not the dorsal HPC was critical. Previous research has reported that in the No Confinement procedure of RAM, rats show a tendency to turn either clockwise or counterclockwise, which helps simplify the task and reduce the memory load^42^. Rats exhibit an egocentric sequential strategy as a highly efficient approach in working memory tasks in which all arms are baited^43,60^. The serial strategy, which involves a sequence of 45° turns, is acquired by animals as they recognize it as the most effective way to solve the task, especially when all arms are open and baited^39,61,62^. With this strategy, rats learn to forage efficiently, obtaining all rewards while traveling the least possible distance and spending the least amount of time. The clockwise serial strategy is quickly acquired when animals transition to a No Confinement procedure^43^. Consistent with this, we found that the use of the egocentric sequential strategy increased immediately on day 1 of the No Confinement procedure.

The entire HPC appears to be important for acquiring an egocentric sequential strategy in a RAM, but the ventral HPC was most critical, showing a significantly lower score with both 6 and 8 open arms trials following its lesion. Previous studies have revealed that the egocentric strategy remains intact in HPC lesioned animals when performing a T-maze task that only requires a simple turning response^63^; instead, when egocentric memory requires a complex sequentially organized behavior, HPC is involved^64,65^. Egocentric responses have been found unexpectedly in the HPC and they would explain its role in episodic memory^66,67^. The prominent involvement of the ventral HPC in acquiring the strategy aligns with findings of increased ventral HPC activation in mice using an egocentric sequential strategy in a starmaze task^31^. These results also support the notion that the HPC plays a key role in temporal order memory—the ability to remember the sequence of body movements needed for an egocentric sequential strategy^68,69^.

### Conclusions and translational implications

We present the first validated rodent model of supraspan memory, demonstrating that hippocampal lesions selectively impair WM once capacity limits are exceeded. Dorsal HPC supports object WM under high load, ventral HPC enables egocentric spatial strategies, and both subregions contribute to supraspan maintenance. This paradigm bridges human and rodent research, providing a powerful tool to investigate the mechanisms of supraspan memory and their disruption in MTL-related disorders. The establishment of a rodent supraspan model has direct translational relevance. Supraspan deficits are well-documented in patients with MTL lesions, epilepsy, and Alzheimer’s disease^3–5^. Our model provides a platform to dissect underlying mechanisms, including network dynamics (e.g., HPC–prefrontal interactions, attractor states), neuromodulatory regulation (dopamine, acetylcholine), and cellular processes (autophagy, plasticity) previously implicated in WM capacity^37^. Future work can leverage this model to test interventions aimed at expanding supraspan capacity, with potential therapeutic relevance for aging and dementia.

## Materials and methods

### Subjects

The subjects were outbred male CD1 adult mice (10-16 weeks old, Charles River, Italy, RRID: rid_000091), weighing 40-50 g before undergoing stereotaxic surgery. They were housed in groups of 3-5 per cage under controlled conditions at 24±1°C and 55±5% relative humidity, with a 12-hour light: 12-hour dark cycle (lights on from 07:30 to 19:30). Testing took place during the light phase. The experiments were conducted in accordance with the European Communities Council directives and Italian laws regarding animal care.

### Surgery

Mice were randomly assigned to three groups: bilateral excitotoxic lesions of the Dorsal HPC (Dorsal lesion), Ventral HPC (Ventral lesion), or PBS-injected group (Control group). The surgical procedure followed the method described in Sannino et al.^11^. Mice were anesthetized via an intraperitoneal (i.p.) injection of Avertin (2,2,2-Tribromoethanol 97%, Sigma-Aldrich SRL, Milano, Italy) at a dose of 20 µl per gram of body weight. They were then secured on a stereotaxic apparatus (David Kopf Instruments). A bilateral injection needle was inserted into the HPC using the following coordinates based on the mouse brain atlas^35^: for dorsal HPC lesions, AP = -1.9 mm, L = ±1.2 mm, D = -1.6 mm from bregma; for ventral HPC lesions, AP = -4.1 mm, L = ±3.0 mm, DV = -3.7 mm from bregma. A µL Hamilton syringe, connected to the needle by plastic tubing, was used to inject 0.3 µL of 20 mg/mL N-Methyl-D-aspartate (NMDA; Sigma-Aldrich, St. Louis, MN) dissolved in 1% phosphate-buffered saline (PBS, pH 7.4) into each side. The control group of animals was injected with only 1% PBS. Behavioral testing began 10-15 days after surgery. In a separate group of animals, a unilateral injection of Fluoro-Gold (Hydroxystilbamidine bis(methanesulfonate), Sigma-Aldrich) was performed using the same surgical method.

### Histology

Following the behavioral testing, all animals were perfused intracardially under Avertin anesthesia. The procedure involved perfusion first with phosphate-buffered saline (PBS, pH 7.4), followed by a buffer containing 4% paraformaldehyde. The brains were then removed and stored in 4% paraformaldehyde for a week before sectioning. Using a vibratome, the brains were sectioned into 50 µm slices, with every second section mounted and stained using the Nissl method.

Images of the stained sections were captured using a Leica microscope (10x objective) and subsequently analyzed. Lesioned areas were defined by regions of cellular loss observed with the Nissl method. Only animals with accurately placed lesions were included in the statistical analysis.

To ensure lesion consistency, specific histological criteria were applied before final data analysis. The dorsal HPC was defined as the region extending from -0.94 mm to -2.30 mm from the bregma, while the ventral HPC was defined as the region extending from -2.46 mm to -3.80 mm from the bregma. The inclusion criteria were as follows:

1. The lesion had to be confined solely to the HPC, with no impact on extra-hippocampal structures.
2. For the ventral lesion group, no lesion or minimal lesion could occur in the more posterior CA3 area of the dorsal HPC.

### Fluorogold labeling

To evaluate the anatomical connections of the dorsal HPC and ventral hippocampus HPC, outbred male CD1 adult mice (10–16 weeks) were used. Groups included mice with dorsal lesions (n = 4) and ventral lesions (n = 4). Fluoro-Gold (Hydroxystilbamidine bis(methanesulfonate), Sigma Aldrich SRL, Milano, Italy), dissolved at 4% in 0.9% saline, was unilaterally injected into either the dorsal HPC (coordinates: AP = -1.9 mm, L = ±1.2 mm, D = -1.6 mm) or the ventral HPC (coordinates: AP = -4.1 mm, L = ±3.0 mm, DV = -3.7 mm). The injection procedure was identical to the NMDA lesion procedure, with a total volume of 0.3 µl of Fluoro-Gold administered per injection site.

Mice were allowed to recover for 9–10 days post-surgery before undergoing transcardiac perfusion. The brains were removed and stored in 4% paraformaldehyde for one week. They were then sectioned into 50 µm slices using a vibratome, and all slices were collected in 24-well plates (8 slices per well) containing 0.002% PBS with 1% sodium azide (Sodium Azide 99%, Sigma Aldrich SRL, Milano, Italy).

For visualization, slices covering the entire brain were mounted and imaged using a Leica fluorescence microscope (10x objective) equipped with a Cy5 excitation filter and a DAPI emission filter, which allowed for detailed examination of Fluoro-Gold staining.

### Experimental design

The animals were tested in different behavioural tasks with the following order: elevated plus maze task (day 11), 6 Different objects recognition task (6-DOT) (day 12), 6 Identical objects task (6-IOT) (day 22), radial arm maze task (confinement and no-confinement procedure (day 23-39).

### Behavioral procedures

#### Elevated plus maze task

Control (n = 10), Dorsal lesion (n = 10), and Ventral lesioned mice (n = 12), largely the same as those used in the Confinement/No Confinement procedure, underwent the elevated plus maze task. The apparatus consisted of two opposing open arms (37 x 9 cm) and two closed arms (37 x 9 cm), all extending from a common central platform (8 x 8 cm). The maze was elevated 50 cm above the floor and was situated in a dimly lit room (100 W light).

For each test, individual animals were placed in the center of the maze, facing an open arm, and were allowed to explore freely for 5 minutes. The test was video-tracked using a camera mounted on the ceiling, connected to a video-tracking system (AnyMaze, Stoelting, USA). The primary measures recorded were the percentage of time spent in the open arms and the percentage of open arm entries.

#### 6-DOT/6-IOT

6-DOT and 6-IOT were performed as previously described^11,16,36–38^ on the following groups, largely composed of the same mice: Control (n=8), Dorsal lesion (n = 8), Ventral lesion (n = 12). The 6-DOT was designed to measure WM under high memory load conditions, requiring the animals to remember a large number of objects over a short time interval.

For the 6-DOT, mice were first isolated in a waiting cage for 15 minutes prior to testing. They were then subjected to a habituation period (T1 phase) of 10 minutes in an empty arena (35 x 47 x 60 cm). The habituation period served to assess motor impairments. After 1 minute back in their waiting cage, the mice underwent the study phase (T2 phase), during which they explored six distinct objects for 10 minutes or until they accumulated a total of 120 seconds of exploration. Exploration was defined as the time when the mouse’s nose was in contact with the object (within 2 cm).

Following a 1-minute intertrial interval (ITI), the animals entered the test phase (T3 phase), during which the objects were replaced with identical copies of the familiar objects and one novel object. To avoid bias, two different new objects were used, and the position of the new object was randomized across animals. The animals were allowed to explore the objects for 5 minutes. Their behavior during this phase was recorded with a video-tracking system (Any-maze, Stoelting, USA) and analyzed by a trained observer.

In the 6-IOT, the mice were exposed to identical copies of the same object. This design increased the number of objects without increasing the memory load, allowing the 6-IOT to serve as a control task for the 6-DOT. The T1 phase was identical to the study phase in the 6-DOT. During the T2 phase, mice explored the objects for 5 minutes or up to 35 seconds of total exploration. In the T3 phase, the objects were replaced with identical copies of the familiar objects and one new object. Mice explored the objects for 5 minutes.

The 6-DOT and 6-IOT tasks were administered 8 days apart to prevent interference. Novel object discrimination was calculated as a percentage of the total object exploration time using the following formula: New object exploration / Exploration of New object + Exploration of all familiar objects.

### Eight arms radial arm maze

#### Confinement and No Confinement procedure

Mice (Control, n = 10; Dorsal lesion, n = 10; Ventral lesion, n = 12) underwent the WMC version of the radial arm maze task, as previously described (Olivito et al., 2016). The protocol included trials performed under two conditions: a Confinement procedure during the first four days to prevent the use of egocentric spatial memory and a No Confinement procedure for the subsequent four days^16^.

The apparatus was a custom-built wooden radial arm maze with a grey-painted floor, elevated 84 cm above the ground. It consisted of eight identical, evenly spaced arms (38 cm long × 8 cm wide × 9 cm high) radiating from a central octagonal platform (19 cm in diameter). Transparent Plexiglas doors (9 cm high) were positioned at the entrance of each arm as needed. The maze was located in a well-lit room surrounded by grey curtains, with four distinct visual extramaze cues placed around it. A hand-crafted confinement box was used during the confinement procedure. The experimenter maintained a consistent position throughout the testing.

A small piece of cocoa cereal (Coco Pops, Kellogg’s) was used as a reward and placed at the distal end of each arm. To motivate performance, mice were placed on a restricted feeding schedule starting one day before testing and maintained throughout the study, keeping their body weight at 80–85% of free-feeding levels. Water was available ad libitum.

The protocol consisted of four phases: a habituation phase, a pretraining (PT) phase, a WMC training phase with confinement (first four days), and a WMC training phase without confinement (subsequent four days). The *habituation phase* lasted for three days, with one trial per day lasting 10 minutes. During this phase, all arms were open, and pieces of cocoa cereal were scattered throughout the maze to promote exploration. On the second day, the confinement box was introduced to familiarize animals with its use. In the *Pretraining phase*, mice underwent nine training trials per day over two days. Each trial involved two open and baited arms, while the other six arms remained closed. The open and baited arms varied between trials on the same day and across days. Mice were confined in the maze center for 5 seconds after exiting an arm. Trials ended when mice visited both open and baited arms or when six minutes elapsed. Arm entry was defined as the mouse placing all four paws beyond the midpoint of the arm. The sequence of arm entries and the time to complete each trial were manually recorded.

The *WMC training phase with the Confinement procedure* lasted for four days and involved nine training trials per day. The working memory load was adjusted by varying the number of open and baited arms across trials: : 3 trials with 3 open and baited arms (5 closed), 3 trials with 6 open and baited arms (2 closed), and 3 trials with 8 open and baited arms. The selection and order of open and baited arms were randomized across trials and days. Trials ended when mice entered all the open and baited arms or when six minutes elapsed. The sequence of arm entries and the time taken to complete each trial were recorded.

After two rest days (during which mice remained on the restricted feeding schedule), the *WMC training phase with the No Confinement procedure* commenced. This phase replicated the WMC training phase with Confinement, with nine trials per day for four days. The number of open and baited arms (3, 6, or 8) varied across trials. Trials ended when mice entered all the open and baited arms or when six minutes elapsed.

The measures analyzed during both the PT phase and WMC training phase (with both confinement and no-confinement procedures) included the mean number of errors made during trials with 3, 6, and 8 open and baited arms. Errors were defined as re-entering previously visited open arms.

During the WMC training phase, we also measured the frequency of each Span (the sequence of correct entries before making an error), normalized to the total number of trials and mice. The values were then categorized into two groups: the sum of frequencies for Spans ranging from 1 to 4, and the sum of frequencies for Spans ranging from 5 to 8. Additionally, we computed an index derived from the difference between the mean number of errors and the mean span for each animal

#### Automatic Scoring of Behavioral Strategies in the Radial-Arm Maze Task

A Radial Arm Maze (RAM) database was developed in collaboration with the bioinformatics core of the Telethon Institute of Genetics and Medicine (TIGEM) to analyze data scored from the radial arm maze task. The web tool, RAM Analysis, automatically collected scores derived from manually entered data. It was implemented using HTML, PHP, and JavaScript, and the loaded and analyzed data were stored in a PostgreSQL database.

The adjacent pattern measured the total number of entries into adjacent arms (45° turns). The alternating pattern quantified the number of entries made in an alternating fashion (skipping one arm to enter the next, 90° turns). The opposite pattern measured the total number of entries into arms located directly opposite each other (180° turns). A random strategy was also defined as an entry into an arm after skipping three arms, resembling a 135° turn. For each of these patterns, 1 point was added for each entry made in the corresponding direction.

The sequential strategy was defined as the longest correct sequence (incorrect entries were excluded) of adjacent arms made by the animal during a trial. It was calculated by adding 1 point for each arm in the correct sequence. For example, if the sequence of entries was 1-2-3-4-5-6-7-8, the score for the sequential strategy would be 8 (1 point for each correctly entered arm).

For a more detailed description of each measure, please refer to the Supplemental Information. Additional measures are also available on the database website, which can be easily accessed at https://ram.tigem.it/index.php.

#### Statistical analysis

All the measures used in the 6-DOT, 6-IOT, Pre-training phase of the RAM and the Elevated Plus maze were analysed with a one-way ANOVA followed by Tukey’s multiple comparison test. For the Radial Arm Maze task, we used a two-way repeated measures ANOVA followed by Tukey’s multiple comparison test to analyse the mean number of errors, the difference (Errors – Span), the mean score of the sequential strategy, the percentage of strategies at each open arms condition, and the mean number of errors across days at each open arms condition. For the percentage of Span frequency, we used Fisher’s exact test followed by Benjamini-Hochberg multiple correction.

## Acknowledgements

This work was supported by DSB Project. AD004.417.001 “New diagnostic and therapeutic biomarkers of degenerative diseases” (FOE 2022), National Recovery and Resilience Plan (NRRP), project “Cascata-MNESYS (PE0000006) - A Multiscale integrated approach to the study of the nervous system in health and disease” (DN. 1553 11.10.2022) to EDL and by he InvAt “Active and Healthy Aging” (FOE 2022) program.

