## Supplementary for "Functional specialization of the hippocampus along its dorso-ventral axis: from span to supraspan in working memory capacity"

### Materials and methods

*RAM database*

RAM database (https://ram.tigem.it/exp2.php) was projected with the aim of automatically extracting behavioural metrics from an extensive dataset generated through an eight arms radial arm maze task. Originally, it was developed for the working memory capacity version of the radial arm maze task in mice, as described in Olivito et al 2016.

The database comprises four primary sections: *Animals, Training, Archive, Statistics*.

In the *Animal* panel, demographic information pertaining to each new subject can be input.

The *Training* section is dedicated to entering entries sequences of each trial/day of the task, open arms (each identified by a number from 1 to 8) and the duration of each trial.

In the *Archive* section, automatic scoring of the following metrics is provided for each subject across various trials of the task:

- *Adjacent* is a metric that calculates the sum of adjacent entries within each entries sequence. It involves adding +1 for every adjacent entry. For example, if the sequence is 3**67**8855**765**5**87**222**234**6, the “Adjacent” value would be 7. All adjacent entries are highlighted in bold by assigning a value of 1 to each of them.
- *Alternating* is a metric that calculates the sum of alternating entries within each entries sequence. It involves adding +1 for every alternate entry. For example, if the sequence is 367885**57**655872223**46**, the "Alternating" value would be 2. All alternating entries are highlighted in bold by assigning a value of 1 to each of them.
- *Opposite* is a metric that calculates the sum of entries in arms located in front of the last visited one. This is achieved by adding +1 for every entry corresponding to an opposite arm. In an 8-arm radial maze, opposite arms are paired as follows: 1-5, 2-6, 3-7, 4-8. For example, if the sequence is 2**2651**63**15**2**7384**, the "Opposite" value would be 5.
- *Sequential* is a metric that calculates the length of the longest sequence of adjacent entries by counting the number of arms included in this sequence. For instance, if the sequence is 32231153**32176**, the "Sequential" value would be 5.
- *SeqMErr* is a metric that calculates the length of the longest sequence of adjacent entries that have not been previously visited, determined by counting the number of arms included in this sequence. For example, if the sequence is **32**231153321**76**, the "SeqMErr" value would be determined accordingly based on the mentioned criteria.

- *Span* is a metric that calculates the number of correct entries made before committing an error. For example, if the sequence is **43872**182117665, the "Span" value would be 6, indicating six correct entries before an error was made.
- *SpanMSeq* is a measure of the number of correct and not sequential entries before committing an error. In the provided examample: **4831**67**2**84217318348175, SpanMSeq would be 5. Sequential here is the sequence 67.
- *First* is a conventional identifier corresponding to the initial arm in the maze. It signifies the arm number associated with the first entry made in a given sequence. For example, if the sequence is **4**83167284217318348175, the "First" value would be 4, indicating that the first entry was made in arm number 4.
- *Perseverative Behavior* is a metric that calculates the total count of re-entries into the arm that was just visited. For example, if the sequence is 367**8855**76**55**87**222**346, the "Perseverative Behavior" value would be 5, indicating that there were 5 re-entries into the arm that was most recently visited.
- In the *Statistics* section, you can find measures related to error position. These metrics draw upon the concepts of primacy and recency, which are commonly used to describe working memory strategies in humans (Glanzer and Cunitz, 1966).
- *First_Err* defines the conventional arm number where the first error is committed. For example, if the sequence is 3678**8**55765587222346, the "First_Err" value would be 8, indicating that the first error occurred in arm number 8.
- *Dist_FirstErr* defines the position of the first error, starting from the last correct visited arm to the first visited arm within the span sequence. Consequently, "Dist" is 1 if the first error occurs in the last correctly visited arm of the span sequence. For instance, if the sequence is 3678**8**55765587222346, the "Dist_FirstErr" value would be 1, indicating that the first error is in the last correctly visited arm of the span sequence.
- *Cnt_1st_AfterErr* defines the number of re-entries into the first visited arm of the entry sequence after the first error. For example, if the sequence is 36788557655872223**3**46, the "Cnt_1st_AfterErr" value would be 1, indicating that there was one re-entry into the first visited arm after the first error.
- *Dist_FirstErr_Norm* defines the position of the first error, measured as "Dist_FirstErr" normalized by the number of possible positions based on the "Span." For example, if the sequence is 3678855765587222346, and "Dist_FirstErr" is 1 with a "Span" of 4, the "Dist_FirstErr_Norm" would be 0.25.

- *Freq_1$^{st}$_AfterErr* defines the number of re-entries into the first visited arm normalized by the total number of uncorrected entries.

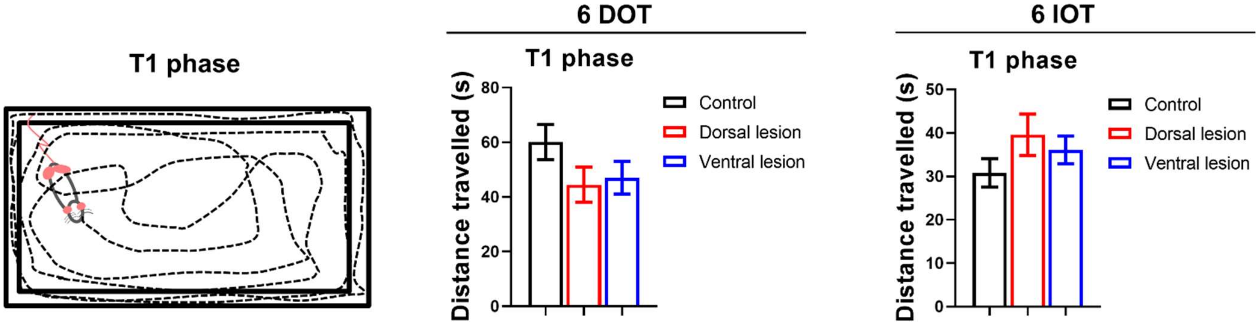


***Figure S1. Effects of dorsal and ventral HPC lesions on locomotor activity.*** *In the T1 phase of the 6-DOT and 6-IOT tasks, mice are free to move within the arena. However, the distance travelled was not affected by either HPC lesion during both the 6-DOT and 6-IOT tasks (6-DOT: lesion* $F_{(2,25)} = 1.571$, $p = 0.228$*; 6 IOT: lesion* $F_{(2,25)} = 1.214$, $p = 0.314$*)*

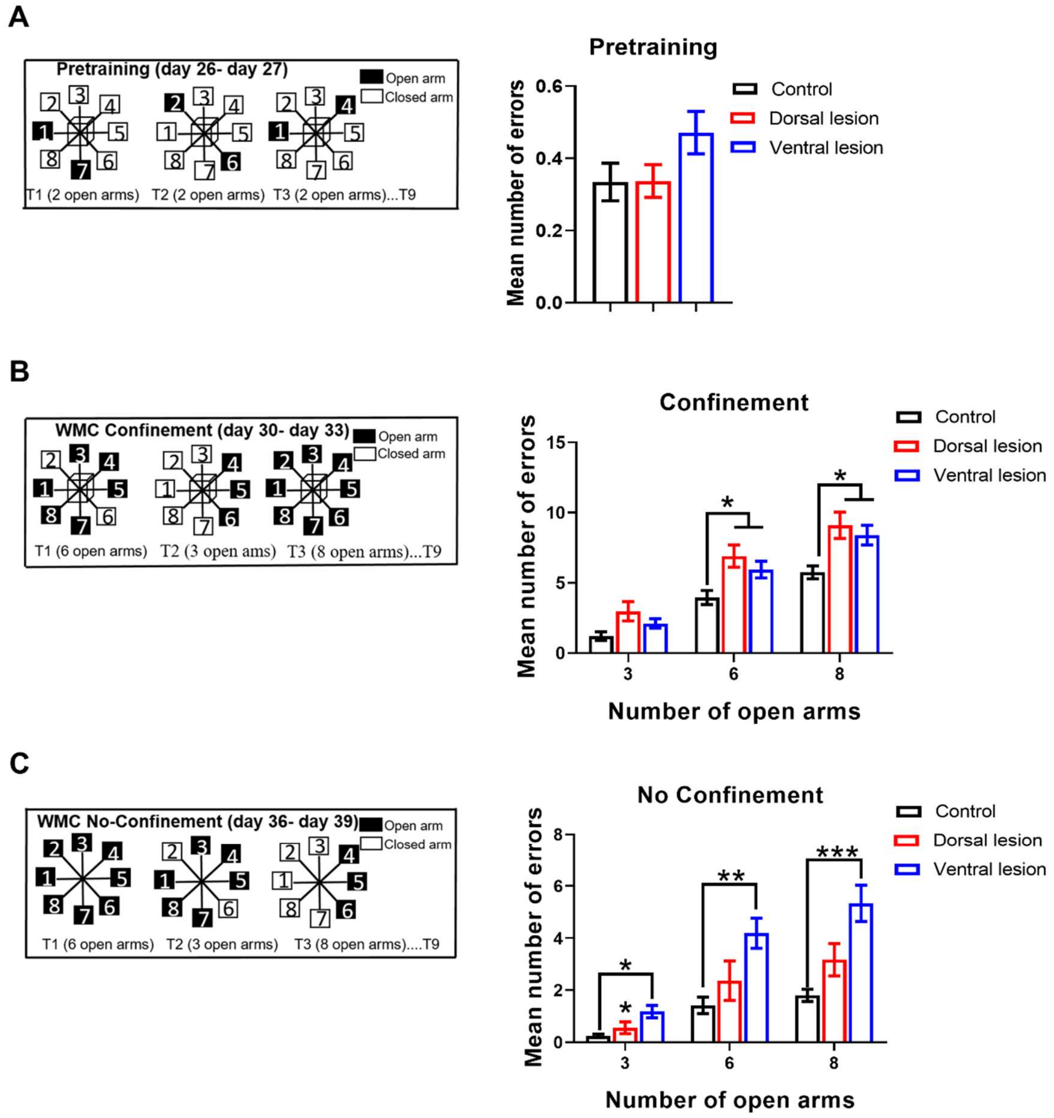


***Figure S2. Mean number of errors across the Pre-training, the WMC Confinement and WMC NO Confinement of the RAM. A)*** *During the pretraining phase, mice learn to avoid re-entering the same arm, as it will no longer be baited. This phase requires them to remember two different spatial locations (arms). Neither HPC lesion affected performance during this phase (one-way ANOVA, lesion* $F_{(2,29)} = 2.262$, *p=0.12).* ***B)*** *In the Confinement procedure, both dorsal and ventral HPC lesion groups made more errors compared to the Control group, especially under the 6 and 8 open arms conditions (two- way ANOVA, lesion* $F_{(2,29)} = 6.843$, *p=0.0037; Tukey's multiple comparisons test, 6 open arms: Dorsal lesion vs. Control p = 0.017, Ventral lesion vs. Control p = 0.048; 8 open arms: Dorsal lesion vs. Control p = 0.017, Ventral lesion vs. Control p = 0.014).* ***C)*** *In the No Confinement procedure, the ventral HPC lesion, but not the dorsal HPC lesion, impaired performance across all memory load conditions (two- way ANOVA, lesion x number of open arms,* $F_{(4, 58)} = 4.761$, *p=0.002; Tukey's multiple comparisons test, 3 open arms: Ventral lesion vs. Control p = 0.006, 6 open arms: Ventral lesion vs. Control p = 0.002, 8 open arms: Ventral lesion vs. Control p = 0.0009) Data are expressed as mean ± SEM*

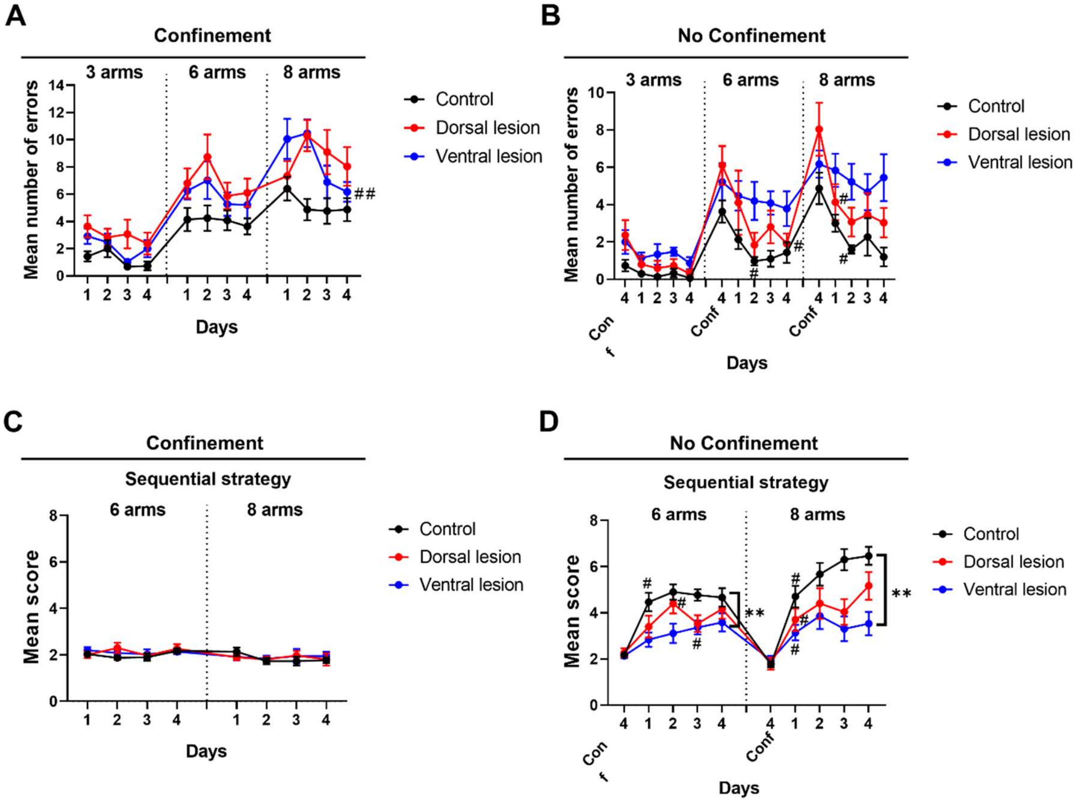


***Figure S3. Acquisition of the radial arm maze task and the egocentric sequential strategy over days following dorsal and ventral HPC lesions. A)*** *A mild effect of HPC lesions on the learning process of the radial arm maze task was observed, specifically under the Confinement procedure with only eight open arms (two-way ANOVA, lesion x days* $F_{(6, 87)} = 2.303$, $p = 0.041$*). Surprisingly, a significant improvement in the performance was observed at this open arms condition upon Ventral HPC lesion (Turkey's multiple comparisons test, day 4 vs. day 1* $p = 0.0092$*).* ***B)*** *In the No Confinement procedure, Control and Dorsal HPC lesioned mice but not Ventral HPC lesioned mice improved their performance over the days compared to the last day of the Confinement procedure (Conf 4).* ***C)****The use of the Sequential strategy was constant across days in the Confinement procedure independently on the lesion.* ***D)*** *In the No Confinement procedure a robust effect of the lesion was observed at both 6 (lesion* $F_{(2, 29)} = 6.569$, $p = 0.0044$*) and 8 open arms condition (lesion* $F_{(2, 29)} = 8.049$, $p = 0.0017$*). Data are expresse as mean ±SEM. #* $p < 0.05$*, ##* $p < 0.01$*, Turkey's multiple comparisons test.*
